# Engram Stability and Maturation During Systems Consolidation Underlies Remote Memory

**DOI:** 10.1101/2022.07.31.502182

**Authors:** Ron Refaeli, Tirzah Kreisel, Maya Groysman, Adar Adamsky, Inbal Goshen

## Abstract

Remote memories play an important role in how we perceive the world and are rooted in ensembles in the CA1 and ACC, however the evolution of these components during systems consolidation has not yet been comprehensively addressed. By applying transgenic approaches for ensemble identification, CLARITY, retro-AAV and rabies virus for circuit mapping, and chemogenetics for functional interrogation, we addressed the dynamics of CA1-ACC ensembles and their connectivity as well as the contribution of astrocytes to the process. We found that the CA1 engrams remain stable between recent and remote recall, and, the inhibition of the engram for recent recall during remote recall functionally impairs memory. We also found that the new cells in the remote recall engram in the CA1 are not added randomly, but differ according to their connections: First, the anterograde CA1 → ACC engram cell projection grows larger. Second, in the retrograde projections, the ACC reduces input to CA1 engram cells, while input from the entorhinal cortex and paraventricular nucleus of the thalamus increases. Finally, we found that activating CA1 astrocytes during acquisition improves recent but not remote recall, and that CA1 → ACC projecting cells are recruited earlier when the astrocytes are stimulated. Our results shine fresh light on systems consolidation by providing a deeper understanding of engram stability and maturation in the transition from recent to remote memory.

## Introduction

Our remote memories, weeks to decades old, define who we are and how we experience the world, however the majority of research in this field has been dedicated to recent memory. Recent memory recall represents synaptic consolidation during the few days following acquisition and has been extensively studied due to its precise time-frame (*1, 2*). Remote memory, on the other hand, is consolidated over much longer periods of time and involves reorganization of multiple brain regions in a process termed systems consolidation (*3, 4*). Even after decades of research, systems consolidation is not fully understood, and the processes underlying the transition from recent to remote memory remain obscure.

The reactivation of the same group of neurons which were originally active during its acquisition is considered the basis of memory recall (*5*). This ensemble of cells, spread throughout the brain, is called an ‘engram’, and reactivating a portion of this sub-population is sufficient to recall a specific memory (*6–9*). Studies even managed to create a false memory by activating engram cells of one context while mice were exploring another (*10*). It has been shown (both theoretically and experimentally) that the engram for a particular memory is distributed across cortical and subcortical brain structures, therefore several brain regions need to be reactivated in order to support a strong recall (*11–13*). When a memory updates, the engram cells supporting it remodel as well (*14, 15*).

In this work, we aimed to follow the evolution of recent and remote memory engrams: to discover whether the remote engram remains stable and whether it is even dependent upon the recent engram on one hand and to determine how it matures with the addition of new cells to the ensemble on the other.

What are the similarities between engrams of the same memory in sequential recalls? Previous studies examined only the overlap between the acquisition engram with recent or remote recall engrams and showed that there is a significant reactivation in hippocampal CA1 as well as in other brain areas (*16–18*). Furthermore, several groups (e.g. (*10, 17, 19–21*) manipulated the activity of the acquisition ensembles to show their necessity and even sufficiency for recent memory, and one study (*17*) observed this in remote memory as well. It is unknown, however, whether the neurons active during remote recall are the same cells active during recent recall, or if both populations partially overlap with the acquisition engram cells but not with one another.

Studying how the engram cell population changes across time requires a method of tagging active neurons more than once during a memory-related task. One approach is targeting immediate early genes, since their transcription occurs when a neuron is hyperactivated (e.g. if it is involved in a specific memory), and because they can be tagged at a specific time window while the animal remains unharmed (*22*). For example, 4-hydroxytamoxifen (4-OHT) generates expression of the desired protein only in neurons expressing the immediate early gene cFos by activating the tamoxifen-dependent recombinase CreER in these cells (*18, 22, 23*). CreER can be expressed virally or in transgenic mice. Since 4-OHT leaves the body within a few hours, the labeling time window is significantly more precise than that of other methods such as the DOX-based system which is approximately a day long (*5, 24–26*). The use of these methods combined with immunohistochemistry (IHC) staining at the second time point in the memory task enables the study of new questions regarding the stability of engrams in reactivation and in the transition between memory stages.

Recalling a memory requires activity elevation in multiple brain structures (*11, 12*). When investigating changes in recent versus remote recall engrams, understanding the dynamics in the long-range connections between different brain regions is essential. Anatomically, the connections between two different brain structures do not alter, but the selection of cells within each structure comprising an engram can certainly change which may be due to their specific projection target. CLARITY (*27, 28*) is a method that enables the clarification of an entire brain, thus allowing one to trace all anterograde or retrograde projections of a given group of neurons (*29*). Anterograde projections of engram output can be observed by labeling cFos expressing neurons with a cytoplasmic fluorophore, while retrograde projections, showing all the cells impinging on the engram population, require a pseudo-rabies that can leap back from the expressing neurons to their pre-synaptic cells (*30*). Thus, the combination of CLARITY and viral-induced methods allows tracking the afferent and efferent projections across the brain regions of interest.

Prominent theories based on human and animal studies suggest that the consolidation of remote memories is a dynamic process requiring hippocampal activation at the beginning(*31, 32*), but relying on frontal cortical regions such as the anterior cingulate cortex (ACC) during more advanced stages (*3, 33–37*). For example, the ACC is involved in the reactivation of remote memories in case of hippocampal malfunction. Recent studies, however, have modified our understanding of remote memory consolidation regarding the ACC and hippocampal time-dependent interactions, suggesting that the hippocampus is needed whenever a memory is recalled, regardless of the length of time passed from memory acquisition (*38*), while the ACC is important at the earliest stages of memory as well as at the remote time point (*39, 40*).

Lastly, the neuronal population can be affected by surrounding astrocytes when a memory task is performed. Each astrocyte in the CA1 engulfs approximately 14 pyramidal cell somata (*41*), envelopes multiple synapses, and can modify activity (*42*). It was shown that astrocytic activation during acquisition of a memory improves recent memory recall (*43, 44*), and astrocytic manipulation during acquisition inhibits remote (but not recent) recall (*45*). However, the involvement of astrocytes in the transition from recent to remote memory was never investigated.

In this study, we marked engram cells at two different time points during contextual fear conditioning in the same mice, and characterized the stability of their reactivation. First, we found that engram neurons are stable (i.e. the ensembles overlap more than expected) between recent and remote recall. Consequently, recall is functionally impaired when the recent recall engram cells in the CA1 are inhibited during remote memory. We then defined changes in the anterograde long-range projections from CA1 engram cells in clarified brains as the memory ages and found that the CA1 → ACC projection strengthens over time. Consistent with this, memory was impaired upon chronic inhibition of the CA1 → ACC projection during the time between recent and remote recall. Next, we targeted all brain structures that provide input to CA1 engram neurons and observed how they change over time. We found structural and functional stability in CA1 engram cells between recent and remote recall and discovered that the new cells do not join the remote ensemble randomly but rather are selected according to their anterograde and retrograde connectivity. Finally, we activated CA1 astrocytes during memory acquisition and observed an improvement in recent memory accompanied by an elevated number of cells projecting to the ACC.

## Results

### Hippocampal CA1 engram cells remain stable from recent to remote memory

Studying the stability of memory engrams requires double labeling of cell activity within the same animal. To tag memory engrams, we injected AAV5-cFos-CreER, inducing the expression of Cre under the promotor of the immediate early gene cFos (that is elevated in active neurons) to the dorsal-CA1 (dCA1) of Ai14-reporter mice (129S6-Gt(ROSA)26Sor^tm14(CAG-tdTomato)Hze^; see methods), conditionally expressing tdTomato in cells that express Cre. The activity of Cre on tdTomato is limited to a 4-8hr time window (*28*) defined by injection of 4-OHT (25 mg/kg, intraperitoneal, i.p.), allowing CRE translocation into the nucleus and consequently tdTomato expression (Figure 1A-B). Three weeks after viral injection, mice underwent fear conditioning training, in which a foot shock was paired with a novel context, and recall was assessed at the recent (2 days) or remote (28 days) time period. At both points, the mice exhibit increased freezing compared to acquisition (F_2,18_=17.65, p=0.000057) (Figure 1C). A separate group of mice did not undergo fear conditioning and were only exposed to their home cage as a control. Animals were injected with 4-OHT 60min after the relevant behavior in order to fluorescently tag all active cells during that task. We found that during memory acquisition and recent or remote recall, 4-OHT introduction caused 16.89, 18.57 and 14.65 percent (respectively) of the CA1 cells to express cFos, while in control mice (which remained in their home cage) only 11.22 percent of the cells were active (F_3,100_=5.543, p=0.002) (Figure S1A). To tag the active cells in two engrams of the same memory in the same animals, the earlier time point was tagged in red (tdTomato) using the 4-OHT to Ai14+cFos-creER as explained above (1^st^ tag, in-vivo), and the later time point was tagged in far-red (Alexa Fluor 647) by IHC against the cFos protein (2^nd^ tag, ex-vivo). Reactivated cells are both tagged in red and stained white (Figure 1D-E).

**Figure 1:**
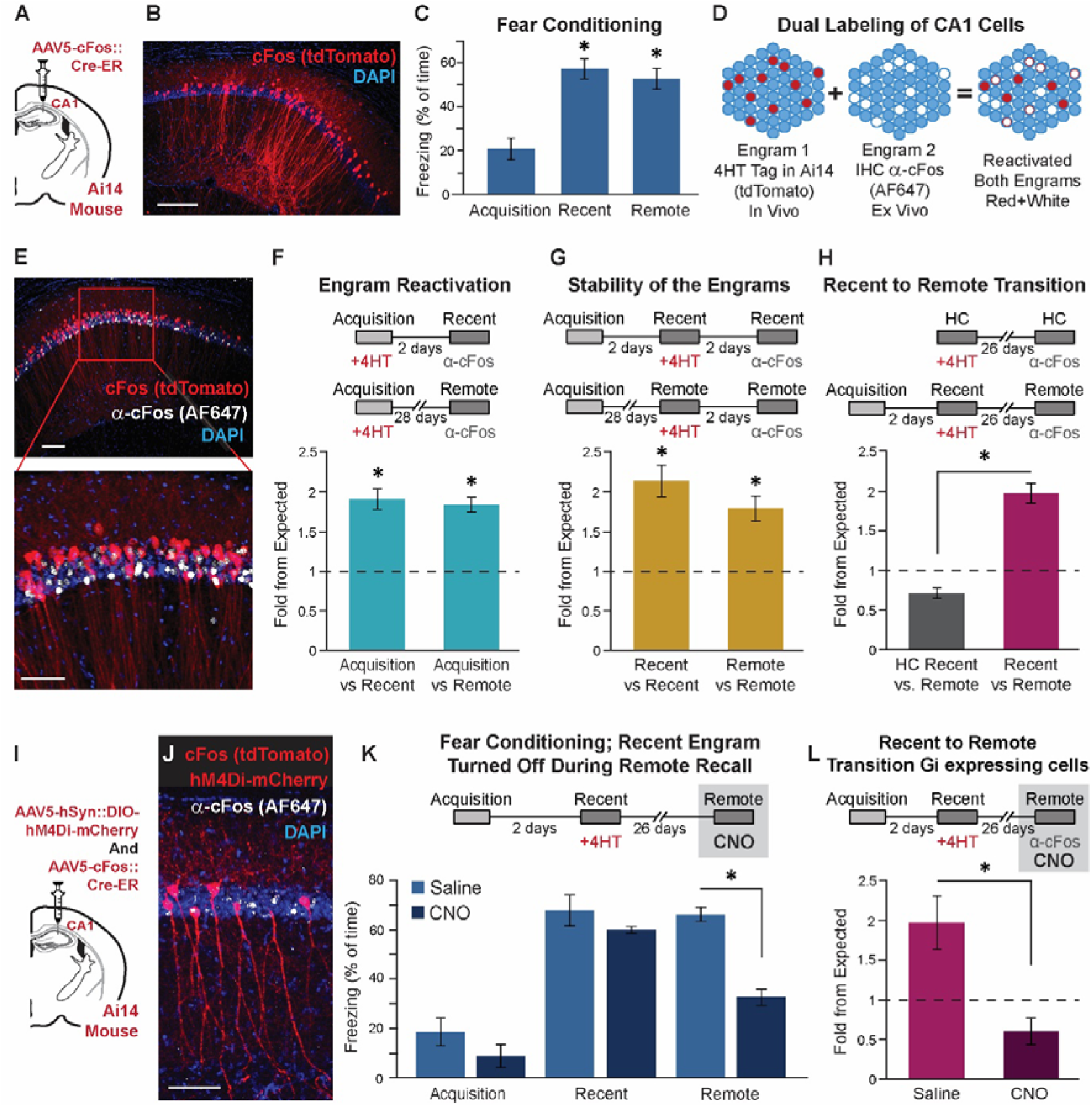
CA1 engram neurons are stable across recent and remote memory transition and the recent ensemble is crucial for remote recall. **(A)** Ai14 mice were injected with a cFos-CreER vector into the dorsal-CA1. **(B)** Cells that were active and thus expressed cFos during 4-OHT administration express tdTomato (red). Scale bar = 100μm **(C)** Mice performance during fear conditioning paradigm (n=7). Freezing is apparent at both the recent and the remote time points (acquisition-recent p=0.0001; acquisition-remote p=0.00043). **(D-E)** Dual labeling of cFos in the CA1. tdTomato (red) in cells from the first time point and α-cFos IHC (AF647, white) in all cFos positive cells during the second time point. Reactivated cells are both red and white. Scale bar = 50μm. **(F)** Reactivation of the acquisition positive cells (tagged first, with 4-OHT) during both recent (n=5) and remote (n=5) recall (tagged second with α-cFos IHC). The acquisition engram cells are significantly reactivated. (p=0.000005 and p=1.725E^-8^). **(G)** Stability of the engram cells two days apart during both recent (n=14) and remote (n=6). There was significant stability at both time points (p=0.000046 and p=0.000636). **(H)** Comparing recent and remote CA1 engram cells. The ensembles are stable during the transition from recent to remote memory (n=7), compared to home caged controls (n=7) (p=5.1551E^-12^). **(I)** Ai14 mice were injected, like before, with a cFos-CreER vector and also with hM4Di-mCherry into the dorsal-CA1. **(J)** Double labeling of cFos and hM4Di-mCherry in the recent engram, and α-cFos after remote recall. Scale bar = 50μm **(K)** Behavioral performance of the hM4Di expressing mice treated with either CNO or Saline (CNO n=5, Saline n=3). Significant reduction in freezing behavior is apparent during remote recall when CNO is applied (p=0.00046). **(L)** Recent to remote transition reactivation is reduced when CNO is applied, compared to saline (p=0.0026). Data presented as mean ± standard error of the mean (SEM).

First, we investigated the reactivation of acquisition CA1 engram cells during recent or remote memory recall. As shown in previous studies (*16, 17*), reactivation of the CA1 acquisition engram exceeded chance levels in both recent and remote recall (see quantification in methods section). Specifically, we found the overlap between acquisition and recent recall to be 190% of expected, and acquisition and remote recall to be 183% (t_(10)_=8.885, p=0.000005, t_(19)_=9.281, p=1.725E^-8^, respectively) with no difference between them (p>0.5)(Figure 1F, S1B). Next, for the first time, we checked the stability of the engram within recent or remote memory twice with two days between each examination. One group of mice was fear conditioned and tested for recent recall two days later and again at day four, while a second group of mice was fear conditioned and tested for remote recall at day 28 and then again at day 30 (Figure S1C). The time interval between the cFos tagging with 4-OHT to the IHC labeling was two days, allowing enough time for the 4-OHT to clear from the body. We found that the stability of engram activation over two days during both recent and remote recalls was high (212% and 178% of chance level, t_(13)_=5.978, p=0.000046 and t_(10)_=4.886, p=0.000636, respectively) (Figure 1G). Finally, we examined, for the first time, whether there exists an overlap between recent and remote engrams, as past research only showed the overlap of each to acquisition individually (*16*). We found that the overlap of cFos labeled cells during recent and remote recall was significantly high (193% of chance level; t-test; t_(19)_=6.9, p=0.000001), as opposed to home cage mice that were tagged at the same intervals without a memory task performance and displayed lower reactivation than expected (t-test; t_(24)_=4.475, p=0.00016). These groups significantly differ from one another (70.7% of chance level; t_(43)_=9.42, p=5.1551E^-12^) (Figure 1H).

The significant overlap between recent and remote engrams raises the question of how turning off the recent engram during remote recall will affect remote memory. To test this question, we injected Ai14 mice with AAV5-hSyn::DIO-hM4Di-mCherry and AAV5-cFos::Cre-ER (Figure 1I), together allowing the induction of hM4Di chemogenetic inhibitor (and tdTomato) only in cells that expressed cFos during 4-OHT injection in recent recall (Figure 1J). Since tdTomato and mCherry are both red, we performed IHC staining against mCherry, and found high penetrance, with >95% of active cells during recent recall (tdTomato) also expressing hM4Di (mCherry)(Figure S1D). 30min before remote recall, we administered CNO (10mg/kg, i.p.) thus inhibiting the recent engram cells prior to the task and preventing their reactivation. Animals treated with CNO during remote recall showed a dramatic impairment in memory retrieval compared to Saline injected controls (t-test; t_(6)_=6.885, p=0.00046) (Figure 1K). The percent of cFos+ cells from the CA1 pyramidal layer cells during recent recall was similar across Saline and CNO groups, but during remote recall (the time of CNO administration), the CNO injected mice showed less cFos+ cells (t-test; t_(14)_=3.44, p=0.004)(Figure S1E). While the overlap of cFos labeling during recent and remote recall was significantly higher in Saline injected mice (196% of chance level; t-test; t_(7)_=2.54 p=0.0388), CNO injected animals showed no difference from chance level (p>0.2), and there was a significant difference between the groups (t-test; t_(14)_=3.66, p=0.0026)(Figure 1L).

Our results support the well-known fact that the CA1 is necessary for memory acquisition and recent recall (*46, 47*) as well as the still controversial notion that the CA1 is involved in remote recall (*38, 48, 49*). Recent and remote recall engrams both involve reactivation of the cells that were active during the acquisition of the memory as previously shown (*16*); they are stable over a two day period; and, most importantly, the engrams overlap during the transition between recent and remote recall. We showed that this overlap is functionally important and that the remote engram relies on the recent engram, since preventing the recent recall engram from reactivation during remote memory impairs recall.

### Imaging of *anterograde* projections from dCA1 points to higher likelihood of reactivation in the CA1→ACC projecting cells as the memory ages

Targeting the anterograde connections of the hippocampal engram neurons at different time points can reveal the dynamic relations with their downstream brain structures. These connections were imaged in whole clear brains. In the first experiment, we quantified the distribution of dCA1 active cell projections throughout the entire brain by tagging the active cells as we did before: Ai14 mice injected with cFos::Cre-ER, and then administered 4-OHT one hour after the time of behavior. Four weeks after tagging the active cells, mice were sacrificed and a CLARITY(*28*) procedure was performed in order to turn the brains transparent, allowing imaging of full hemispheres with single axon resolution (Figure 2A, Movie 1). The dCA1 projects to several targets; the largest projection reaches the mammillary bodies (MB), the second reaches the nucleus accumbens (NAc), and the smallest projections reach the ACC (Figure 2B-F). For all behavioral groups of mice, we found the same total number of axons projecting from the dCA1 engram cells (Figure S2A). We found no significant changes in the number of axons or the percent of axons from total throughout the memory progression in the MB (p>0.661) or the NAc (p>0.664) (Figure 2G-H, S2B-C). The ACC paints a different picture entirely: in home cage, acquisition, and recent recall, only a small portion of the axons from cFos-positive cells head toward the ACC, while during remote recall, the portion of active cells in the CA1 which send their axons toward the ACC doubles (F_3,13_=5.42, p = 0.012) (Figure 2I, S2D).

To further probe our observations, we injected a retrograde virus (AAVretro-CaMKII-eGFP) into the ACC to target the sub-population of neurons within the dCA1 which send their projections toward the ACC (CA1→ ACC) and counted all infected cells in the CA1. In the same mice, cFos expression (tdTomato) during recent or remote recall was labeled as before (Figure 2J-K), and the mice underwent fear conditioning (Figure S2E). The likelihood of activation in the sub population of CA1→ ACC increased as the memory aged (F_3,121_=9.143, p=0.000017)(Figure 2L). However, even at its highest, it reached only the expected value in remote recall, and was significantly lower than expected for home cage, acquisition, and recent recall.

**Figure 2:**
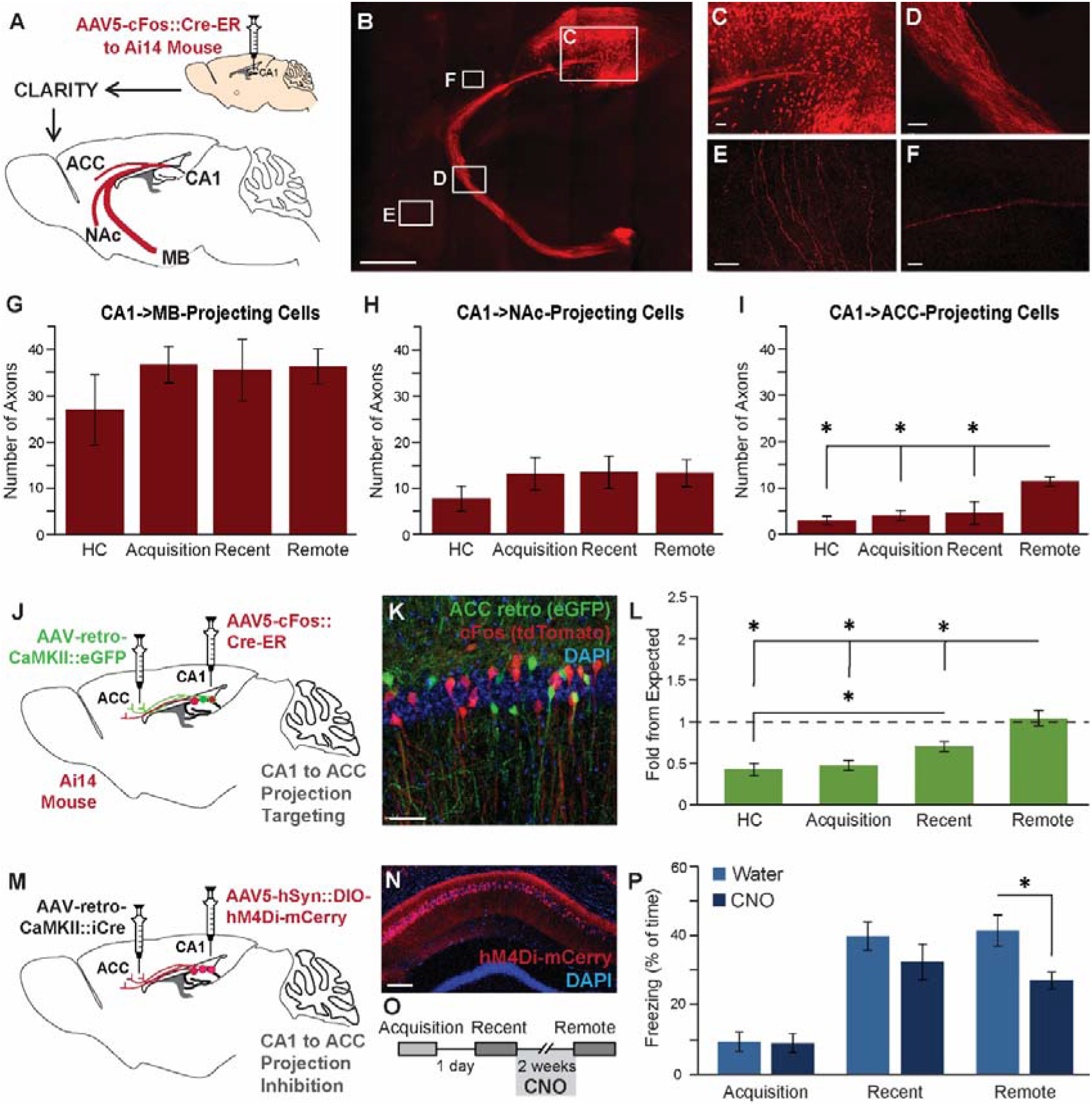
CA1→ACC projecting cells recruitment increases with memory maturation. **(A)** Ai14 mice were injected with a cFos-CreER vector into the CA1 to tag active cells. CLARITY procedure allowing full brain imaging was performed at the different memory stages. **(B)** An entire hemisphere of a mouse expressing tdTomato (red) in cFos positive cells. Scale bar = 1mm **(C-F)** Zoomed-in images of the CA1 engram cell bodies. Scale bars = 10μm **(C)** the fornix going toward the MB **(D)** the axons projecting toward the NAc **(E)** and the projection toward the ACC **(F)**. **(G-I)** The number of tdTomato positive axons counted at the different memory stages going toward the MB (p=0.661). **(G)**, the NAc (p=0.664) **(H)** or the ACC **(I).** Only in this region, the remote recall projection size is double than that of the other time points (HC-remote p=0.026; acquisition-remote p=0.033; recent-remote p=0.039). **(J)** Ai14 mice were injected with a cFos-CreER vector to tag active cells in the CA1 and AAV-retro-CaMKII::eGFP into the ACC to tag cells in CA1 projecting to the ACC (CA1→ACC). **(K)** CA1 → ACC cells express eGFP (green) and the active cFos positive cells express tdTomato (red). Scale bar = 50μm **(L)** The level of CA1 → ACC neurons participating in the engram increases as the memory ages (HC-recent p=0.034; HC-remote p=0.000074; acquisition-remote p=0.000125; recent-remote p=0.021). **(M)** AAV-retro-CaMKII::iCre was injected in the ACC, and the Cre dependent hM4Di-mCherry virus was injected into the CA1, and tagged only CA1 →ACC neurons. **(N)** hM4Di (mCherry) CA1 →ACC cells in the CA1. Scale bar = 100μm **(O)** Behavioral paradigm. Mice were continuously exposed to CNO between recent and remote recall to inhibit CA1→ACC cells during systems consolidation. **(P)** Memory performance was impaired in the CNO group (n=7) where the CA1 ACC cells were chronically inhibited, compared to the water control group (n=8)(p=0.018). Data presented as mean ± SEM.

We previously showed that inhibiting the CA1 → ACC projection during acquisition impairs remote memory (*45*). To check the functional significance of the CA1→ACC projection in the transition from recent to remote memory, we injected wild type mice with AAVretro:CaMKII-iCre in the ACC, and AAV5-hSyn:DIO-hM4Di-mCherry in the CA1 thus enabling specific inhibition of the CAH→ACC neurons (Figure 2M-N). The mice underwent fear conditioning acquisition, performed recent memory recall, and were then administered CNO via their drinking water for two weeks (10mg/kg/day) before remote recall (Figure 2O). Remote recall was reduced in the CNO group compared to the controls that were received water (t-test; t_13_=2.7, p=0.018) (Figure 2P).

Lastly, we labeled dCA1→NAc cells in a different group of mice by infecting the NAc with the same retrograde virus (Figure S2F-G). The mice underwent fear conditioning (Figure S2H), and the likelihood of activation within the sub population of CA1→NAc projecting cells did not differ between recent and remote recall (Figure S2I).

Our results in this section reveal that the projection of dCA1 engram cells to the ACC increases with the transition from recent to remote recall. Furthermore, the cells constituting this projection become increasingly involved in the engram. Finally, we showed that CA1→ACC cells are functionally involved, as remote memory recall is impaired if the projection is inhibited during systems consolidation between recent and remote recall.

### Rabies based *retrograde* projections from dCA1 changes as the memory ages

The hippocampus receives and integrates input from multiple brain structures. To investigate the input sources of the CA1 ensemble cells, i.e., which cells impinge upon them during the different stages of memory acquisition and recall, we used the pseudorabies approach (*30*) that was used for a different purpose in the CA1 (*50*) and was recently used to study recent memory in the amygdala (*51*): we first expressed the mutated avian tumor virus (TVA) receptor (TC66T) and glycoprotein (oPBG) in ensemble cells by injecting AAV2-CAG-flex-TC66T-mCherry and AAV2-CAG-flex-oPBG into the dCA1 of TRAP2 mice (Fos^tm2.1(icre/ERT2)Luo^) expressing CreER protein under the cFos promoter. Three weeks later, mice underwent a fear conditioning task, and 4-OHT was introduced during either acquisition, recent, or remote recall, allowing the Cre to enter the nucleus in the active ensemble cells and induce the expression of TVA and the glycoprotein (Figure 3A). Two weeks later, a pseudo-rabies virus (ENV-Rb-ΔG-eYFP) was injected into the same location in the hippocampus, and it infected and complemented only in the cells expressing TC66T + GP, i.e. those which are a part of the ensemble cells (Figure 3B-C), and spread to their presynaptic neurons. After another week, the mice were sacrificed and the brains clarified.

**Figure 3:**
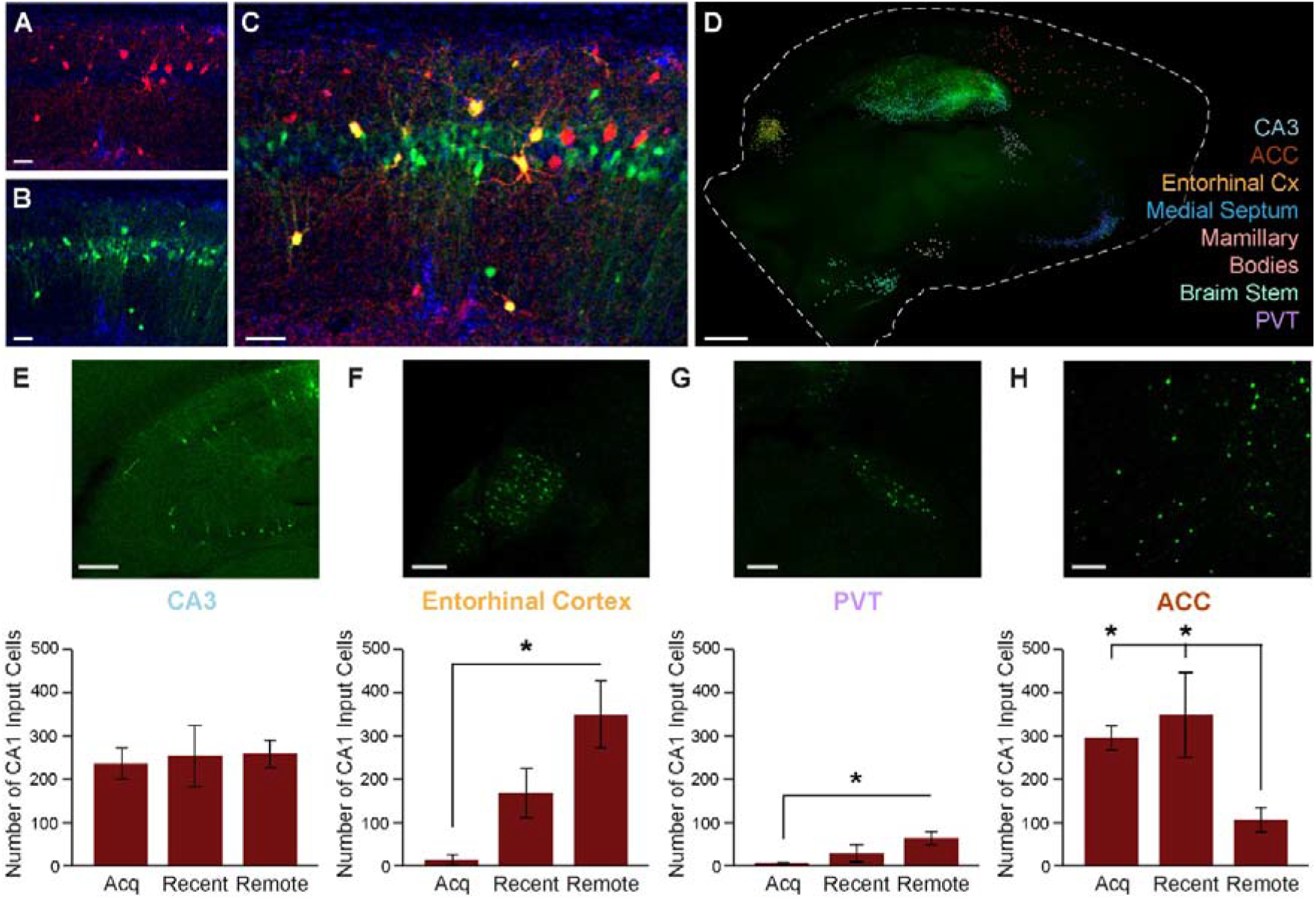
CA1 engram cells receive different brain-wide input as the memory ages. **(A)** CA1 engram cells express TVA-mCherry (red). Scale bar = 50μm **(B)** CA1 cells infected with Pseudo-Rb vector express YFP (green). Scale bar = 50μm **(C)** Overlay of A and B, showing cells that express both mCherry and YFP (43% of mCherry+ cells), are considered starting cells. Scale bar = 50μm **(D)** Whole brain imaging reveals seven brain structures with input cells to the CA1 engram cells. Scale bar = 1mm **(E-H)** Top: input cells in TRAP mice cleared brains. Bottom: number of input cells counted during the different memory stages (acquisition n=7; recent n=4; remote n=5). Scale bars = 100μm. The number stayed the same in the CA3 (p=0.92) **(E)**, increased in the entorhinal cortex (acquisition-remote p=0.0006) **(F)** and the PVT (acquisition-remote p=0.007) **(G)**, and reduced in the ACC (acquisition-remote p=0.007; recent-remote p=0.016) **(H)** as the memory matures. Data presented as mean ± SEM.

We then imaged and analyzed the spatial distribution of the presynaptic cells throughout the entire brain (Figure 3D, Movie 2) at the acquisition, recent, and remote recall time points (fear condition: t(12)=16.44, p=0.00028; t(6)=4.72, p=0.0033 and t(8)=7.27, p=0.000086, respectively) (Figure S3A-C). We located seven different main brain structures that send their projections toward the dCA1 engram cells and counted the number of cells at each memory stage. Notably, we found that the total number of input cells to the CA1 engram remains similar across acquisition, recent, and remote recall (1298, 1273, and 1200 input cells, respectively) (Figure S3D). As expected, CA3, a main projection to the CA1, did not change the number of its input cells into the CA1 engram over time (Figure 3E). The lateral and medial entorhinal cortices and the PVT increase their number of input cells to dCA1 during remote recall (F_2,13_=13.02, p=0.0008; F_2,13_=6.9, p=0.009, respectively) (Figure 3F,G). On the other hand, the ACC decreases the number of input cells into the CA1 during remote recall (F_2,13_=6.33, p=0.012) (Figure 3H). In the medial septum, during remote recall the number of input cells projecting to the CA1 engram decreases, but no difference was found between recent and remote recall (F_2,13_=4.14, p=0.041) (Figure S3E), and no change in the number of input cells at different time points in either the MB or brain stem were observed (Figure S3F,G).

These results provide a comprehensive view of the CA1 engram brain-wide connectivity at different memory stages. It seems that the addition of new cells to the remote ensemble in the CA1 is positively biased in favor of cells with input from the entorhinal cortices and the PVT as opposed to cells receiving input from the ACC.

### Activation of the Gq pathway in CA1 astrocytes during memory acquisition alters CA1 → ACC connectivity and enhances memory performance

Previous studies demonstrated that activation of the Gq pathway in CA1 astrocytes during acquisition of a memory enhances recent recall (*43, 44*) but its role in remote recall is still unknown. Astrocytes were also shown to affect specific projections (*45, 52, 53*), and since we found that the CA1 → ACC recruitment increased over time, we wanted to examine whether astrocytic activation (with hM3Dq) will affect this process. To that end, we used Ai14 mice injected with AAVretro-CaMKII::eGFP into the ACC (tagging the CA1→ ACC projection neurons in green), AAV5-cFos::CreER into the CA1 (tagging all recent recall neurons in tdTomato red), and finally, AAV1-GFAP::hM3Dq(Gq)-mCherry into the CA1, causing expression of hM3Dq in astrocytes (Figure 4A). 4-OHT was administered during recent recall to tag the engram, and the remote engram was stained by IHC against cFos (Figure 4B). Astrocytes expressing hM3Dq-mCherry and recent recall engram cells expressing tdTomato are both red, therefore we stained the astrocytes using an α-mCherry antibody and found that >99% of astrocytes were stained and only <8% of the neurons (Figure S4A). Animals were trained in contextual fear conditioning, and Saline or CNO (3mg/kg, i.p.) was injected 30 minutes before acquisition. As expected (*44*), no behavioral differences were observed during acquisition, but memory performance during recent recall was elevated among the mice treated two days prior with CNO compared to the Saline treated controls (t-test; t_(15)_=3.741, p=0.002) (Figure 4C). During remote recall, no significant difference between the groups was observed. When observing CA1 → ACC projecting cells, we found that Gq pathway recruitment in astrocytes during acquisition resulted in increased activation of this projection during recent but not during remote recall (t-test; t_(25)_=2.421, p=0.023)(Figure 4D).

**Figure 4:**
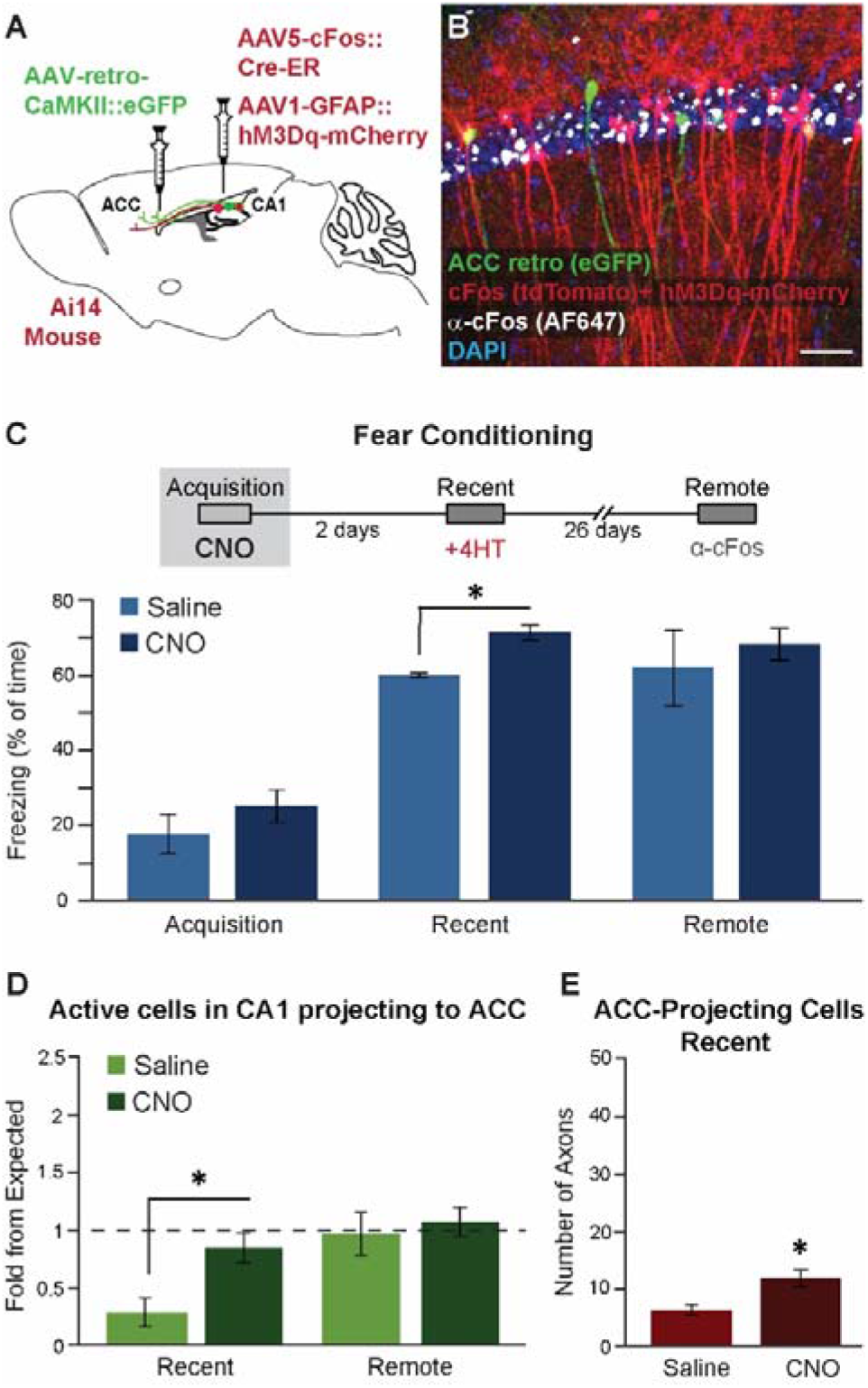
CA1 astrocytic activation during acquisition enhances recent memory and causes changes in the CA1→ACC projecting cells. **(A)** Ai14 mice were injected with a cFos-CreER vector and GFAP-hM3Dq-mCherry into the CA1 to tag active cells and express hM3Dq in astrocytes. AAV-retro-CaMKII::eGFP was injected into the ACC to tag cells in CA1 projecting to the ACC (CA1→ACC). **(B)** CA1 →ACC cells express eGFP (green), hM3Dq positive astrocytes express mCherry (red), recent recall cFos positive cells express tdTomato (red), and remote recall cFos positive cells were stained (white). Scale bar = 50μm. **(C)** Mice were administered saline (n=5) or CNO (n=12) during acquisition. Memory enhancement was observed during recent recall (p=0.0033) but not during remote recall (p=0.38). **(D)** CNO, compared to the saline control group, causes significant recruitment of the CA1 ACC neurons during recent (p=0.023), but not during remote recall (p=0.685). **(E)** The number of tdTomato positive axons of ensemble cells heading toward the ACC significantly increased when CNO (n=3) was introduced during memory acquisition to mice expressing hM3Dq in CA1 astrocytes, compared to the Saline control group (n=6) (p=0.045). Data presented as mean ± SEM.

We next questioned whether the earlier recruitment of CA1 → ACC would be apparent in the number of axons extending toward the ACC from the CA1 and whether it will be specific. To that end, another group of Ai14 mice was injected with the AAV5-cFos::CreER and AAV1-GFAP::hM3Dq(Gq)-mCherry vectors to the CA1 and underwent fear conditioning with Saline or CNO injection during acquisition. Yet again, memory performance during recent recall was elevated among the mice that were treated with CNO compared to the Saline treated controls (t-test; t(4)=6.27, p=0.0033)(Figure S4B). During recent recall, all active cells were tagged by 4-OHT administration. Four weeks later, the mice were sacrificed, the brains clarified, and anterograde axons from the CA1 engram cells were counted. We found that the CNO-injected mice, i.e. those with recruited Gq pathway in astrocytes, showed a higher number of CA1 axons projecting to the ACC compared to the Saline group (t-test; t_(7)_=2.434, p=0.045)(Figure 4E) which was not observed in NAc and MB projecting axons (Figure S4C,D).

These results show that the cognitive enhancement effect of astrocytes comes hand in hand with the anatomical effect on the engram projection, specifically influencing the CA1 →ACC projection.

## Discussion

In this study, we found that CA1 recent recall engram populations remain stable during the transition to remote recall. During the transition from recent to remote recall, the CA1 increases its input to the ACC and, concomitantly, the CA1 → ACC projecting cells become more involved in the engram. The ACC decreases its input, and the entorhinal cortices and the PVT increase their input to the CA1 engram as time progresses. Lastly, we found that activation of CA1 astrocytes with hM3Dq during acquisition of a memory results in improved recent memory performance and in strengthening of the CA1 → ACC projection size and involvement in the engram at this time.

Memory recall depends upon the reactivation of the same group of neurons that were active during acquisition (*7*). In this study, we targeted the relation between recall engrams at different memory stages. One could argue that both recent and remote recall are bound to the acquisition ensemble group as was previously shown (*16, 17*), but not necessarily resembling one another. We found that the CA1 engram population remains stable across recent and remote memory, and by inhibiting the neurons that were part of the recent recall engram during remote recall, we demonstrated that its reactivation is functionally crucial for remote memory. A few past experiments tagged and manipulated recent engrams, and showed that they are sufficient to retrieve memory (*10, 20*). On the other hand, when an engram of one environment is activated while another is learned, it will interfere with the new memory (*54*). Similarly, activating the fear extinction engram will impair memory and inhibiting it will improve memory (*55*). One study showed that activating the acquisition ensemble in mPFC enhances remote memory while inhibiting it impairs remote recall (*56*). Lastly, past experiments have targeted the remote engram and showed that it enhances extinction when activated, and reduces extinction when inactivated (*57*). The relationship between recent and remote engrams was never assessed, to our knowledge, and we show for the first time that inhibiting the recent ensemble damages remote recall.

Remote memories are harder to extinguish than recent ones (*58*), and the similarity between ensembles of two memories is higher when they are formed hours apart compared to a week apart (*59*), therefore we hypothesize that this memory permanency is based on increased ensemble stability. Conversely, we found that both recent and remote engrams in the CA1 are stable to a similar degree when measured two days apart. What is the difference then, if not stability? The CA1 engram remains relatively similar between recent and remote memory, but new neurons still join the remote ensemble. Our findings of changed anterograde and retrograde projections can only stem from the new cells that join the remote engram and constitute the changes from the recent engram.

While investigating the connectivity-based properties of the CA1 engram, we found alterations to its input-output connections. The bi-directional circuit between the CA1 and the MB is essential for memory (*60*), but based on our findings, the size of input from CA1 engram cells to the MB and from the MB to CA1 engram cells is constant. The CA1-ACC circuitry, however, changes as the memory ages. First, the pseudo-rabies experiment shows that fewer ACC cells project to CA1 engram cells during remote memory compared to recent. This result fits the repeated finding that the importance of the ACC grows while the role of the CA1 diminishes in remote memory, and is supported by the wide range of studies showing that the ACC can mediate remote recall even if the hippocampus is inactivated (*3, 34, 35, 61*). Second, more CA1 engram cells send input towards the ACC as the memory ages. This implies that new neurons joining the remote engram are more likely to be ACC-projecting neurons. This does not indicate that the overall communication between the cells of two structures increases, only that a greater portion of these cells take part in the engram. Recent studies have shown the significance of the hippocampus not only during recent recall, but after systems consolidation as well (*38*), which sits well with its increased connection to the ACC.

Two more changes in the retrograde connection, i.e. in the information sources for the CA1, were found. First, more cells from the entorhinal cortices, the major cortical input to the hippocampus, project to the CA1 engram as the memory ages. This brain region is known to play a role in both recent and remote memory (*62–65*), and now we show that it changes during the transition from one to the other. Second, the PVT, a stress-related nucleus known to affect auditory cued memory (*66, 67*), also increases its input toward CA1 engram cells as the memory ages. The selection of new CA1 engram cells is biased positively toward their incoming projections from the entorhinal and PVT and negatively toward their ACC afferents.

Neurons are in tight synergy with astrocytes (*42*), and it has been shown in recent years that specific manipulation of astrocytes can modify recent and remote memory (*43–45*). When we activated CA1 astrocytes via the Gq pathway (with hM3Dq) during memory acquisition, we found improved memory during recent recall (as we have shown before (*44*) but not during remote recall. In addition, we found that the CA1 → ACC connectivity alters: First, as early as recent recall, CA1 → ACC projecting engram cells in astrocyte-activated mice reach the level of recruitment of the remote time point observed in non-treated mice. Second, the recent recall engram in the CA1 sends more axons to the ACC when astrocytes are activated during acquisition. We have previously demonstrated that manipulating the Gi pathways (with hM4Di) in CA1 astrocytes can specifically modulate the CA1 → ACC projection and harm remote memory (*45*), indicating that CA1 astrocytes can bear influence on memory at the resolution of a singular projection. However, it is impossible to say at this point in time whether the increase in the CA1→ ACC projection portion of the recent engram mediates the enhanced memory, or vice versa.

In this study, we present a comprehensive view of engrams in the different stages of memory. We show that the engram population in the dCA1 remains stable across recent and remote recall and establish that it is functionally critical for remote memory. We then demonstrate that the addition of cells to the engram is biased regarding anterograde and retrograde connections. Finally, we activated CA1 astrocytes resulting in an increase in CA1 → ACC connectivity among the engram cells and enhanced memory during recent recall. These findings shed light on the transition from recent to remote memory and the remote ensemble selection mechanisms and bring us closer to understanding the process of systems consolidation.

## Supplementary Material

**Figure S1:**
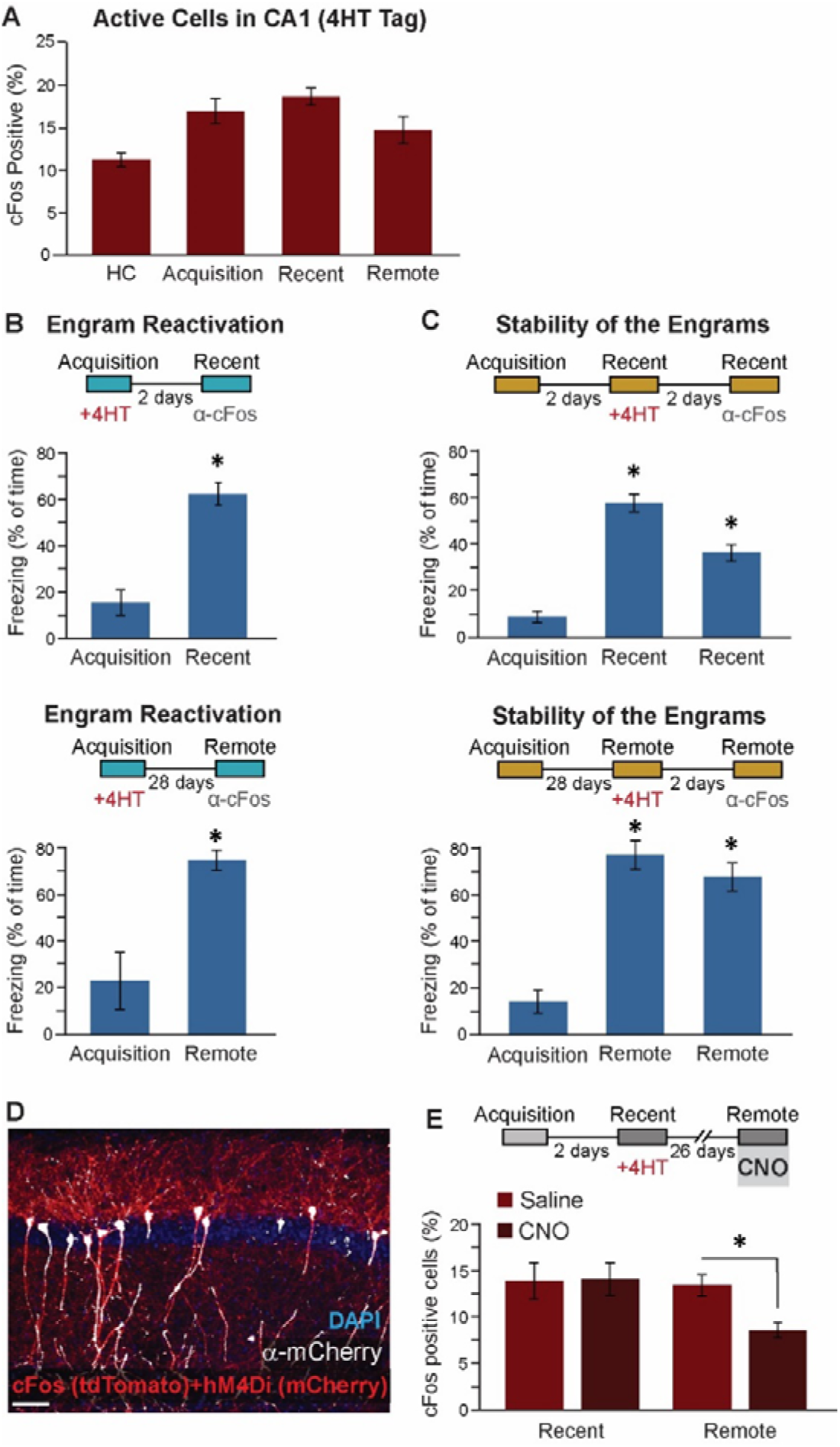
Behavior is persistent throughout experiments, and CA1 recent engram neurons are crucial for remote recall. **(A)** Percentage of cFos positive cells counted at the different memory stages. (**B**) Mice performance during fear conditioning in the engram reactivation experiment. Top: Freezing during acquisition and recent recall (n=5) (p=0.000005). Bottom: Freezing during acquisition and remote recall (n=5) (p=1.725E^-8^). **(C)** Mice memory while exploring stability within memory recall categories. Top: Freezing was measured two and four days after acquisition (n=14) (acquisition-recent p=5.1E^-9^; acquisition-remote p=3E^-6^; recent-remote p=0.0002). Bottom: Freezing was measured 28 and 30 days following acquisition (n=6) (acquisition-recent=4E^-6^; acquisition-remote p=2.4E^-5^). **(D)** Double labeling of red cells (Dio-hM4Di-mCherry and cFos tdTomato) with α-mCherry (156 cells out of 162 cells are both red expressing m-Cherry, indicating >95% penetrance). Scale bar = 50μm. **(E)** Level of cFos positive cells in the CNO group (n=5) and in the saline control (n=3), during recent and remote recall. During remote recall a significant decrease in the level of cFos expression was measured in the CNO group (p=0.004). Data presented as mean ± SEM.

**S2:**
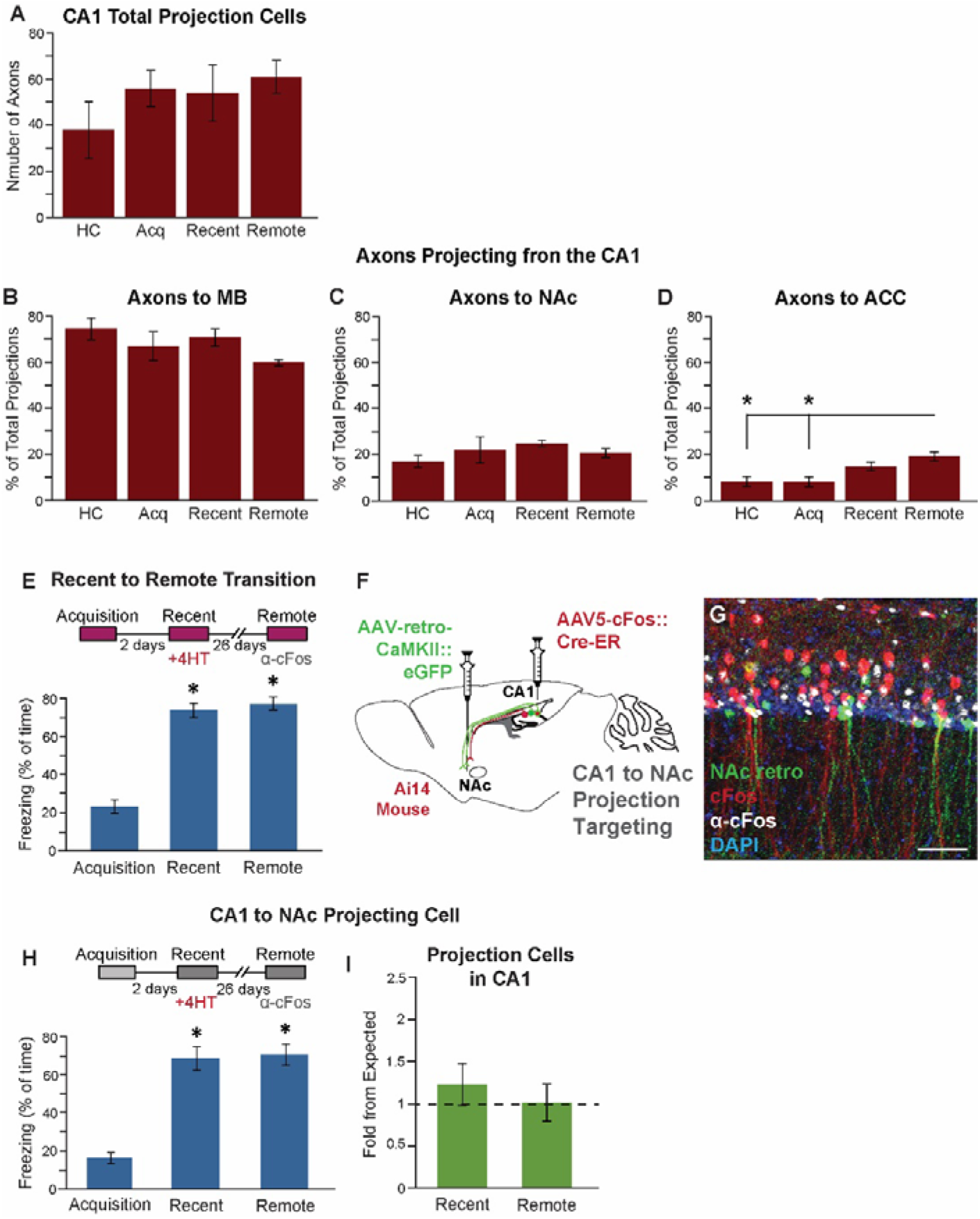
CA1→ACC projecting cells are recruited when memory matures. **(A)** Total number of CA1 projecting engram cells axons counted in clear hemispheres during different time points in the memory maturation. **(B-D)** Percentage of the CA1→MB (p=0.16) **(B)**, CA1→NAc (p=0.46) **(C)** or CA1→ACC axons **(D)**, from the total amount of counted axons, during memory maturation. Only toward the ACC the remote recall projection percentage significantly increased (HC-remote p=0.009; acquisition-remote p=0.004). **(E-J)** Percentage of the active ACC projecting cells out of the total amount of engram cells in the origin areas: entorhinal cortices (p=0.00589) **(E)**, amygdala (p=0.68) **(F)**, PVT (p=0.039) **(G)**, lateral hypothalamus (p=0.82) **(H)**, claustrum (p=0.89) **(I)** and MB (p=0.59) **(J)** during the memory transition from recent to remote. **(K)** Ai14 mice were injected with a cFos-CreER vector to tag active cells in the CA1 and AAV-retro-CaMKII::eGFP into the NAc to tag cells in the CA1 projecting to the NAc (CA1→ NAc). **(L)** CA1→ NAc cells express eGFP (green), the active cFos positive cells during recent recall express tdTomato (red), and all active cells during remote recall are stained by IHC for cFos (white). Scale bar = 50μm. **(M)** Behavioral performance of the CA1→NAc group of mice (n=6) (acquisition-recent p=8E^-6^; acquisition-remote p=5E^-6^) **(N)** The level of CA1 →NAc neurons participating in the engram does not alter from expected during recent recall (p=0.72) nor remote recall (p=0.84).

**Figure S3:**
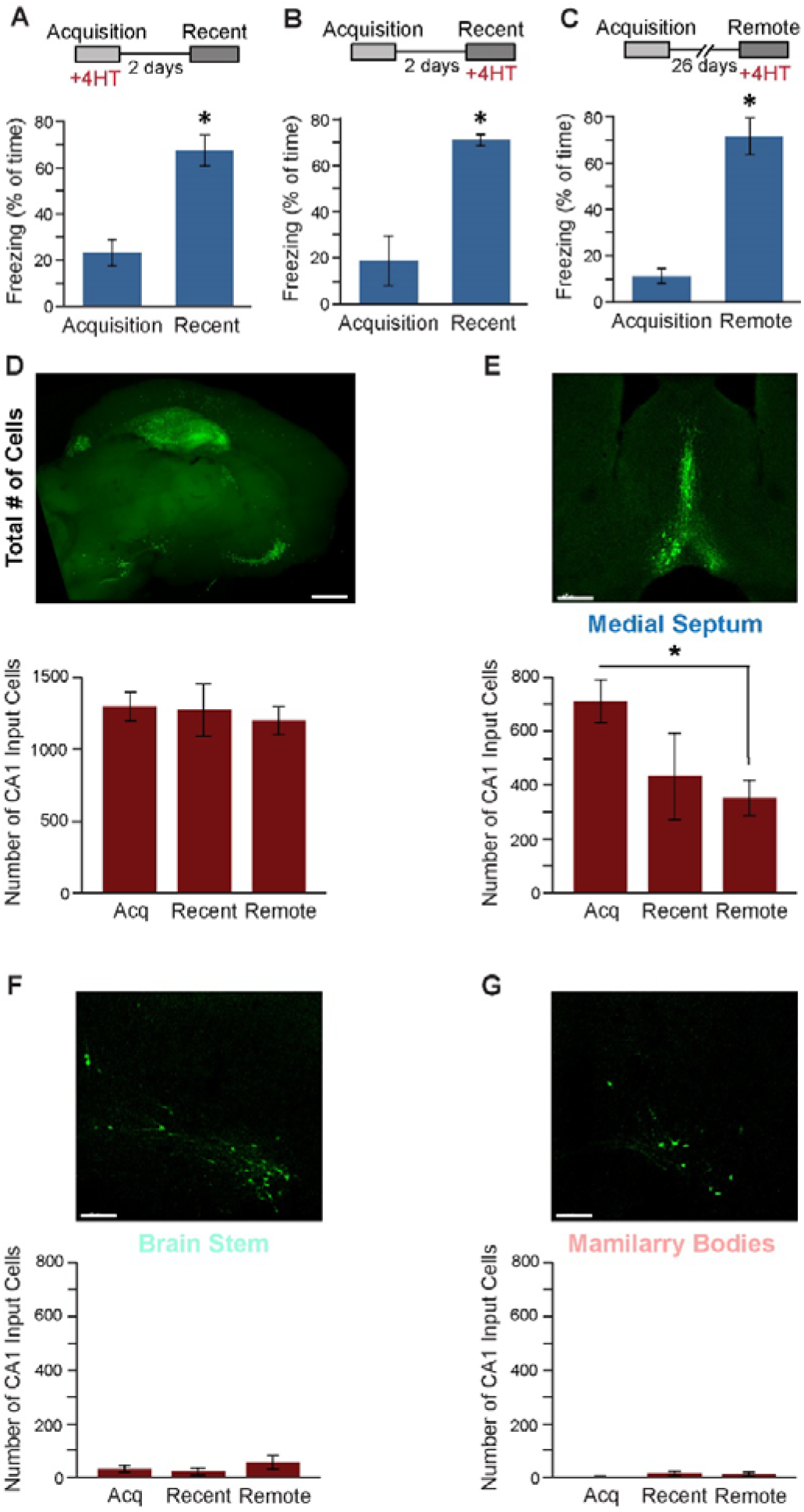
CA1 engram cells receive different brain-wide input during the memory maturation process. **(A-C)** Mice performance during fear conditioning paradigm for CLARITY experiments. Freezing is apparent for the acquisition-tagged mice (n=7, p=0.00028) **(A)**, for the recent-tagged mice (n=4, p=0.0033) **(B)** and for the remote-tagged mice (n=5, p=0.000086) **(C)**. **(D)** Total amount of counted input cells in cleared hemispheres during the memory maturation (p=0.86). Scale bar = 1mm. **(E-G)** Top: Input cells in TRAP mice cleared brains at different brain regions. Scale bars = 100μm. Bottom: Number of input cells counted during the different memory stages in the MS (acquisition-remote p=0.045) **(E)**, in the brain stem (p=0.408) **(F)** and the MB (p=0.24) **(G)** along the memory maturation. Data presented as mean ± SEM.

**Figure S4:**
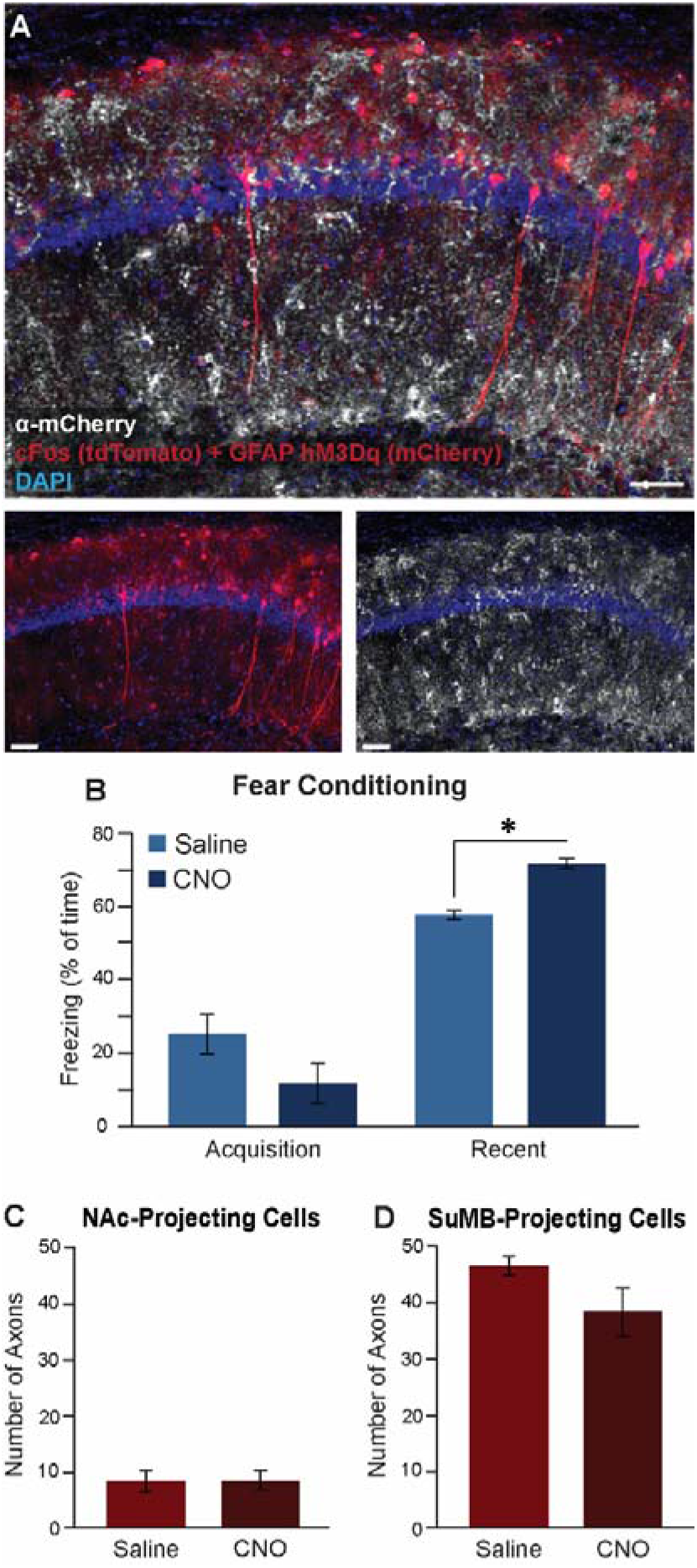
CA1 astrocytic activation does not causes changes in the CA1→NAc nor the CA1→MB projecting cells. **(A)** cFos positive cells express tdTomato (red), hM3Dq positive astrocytes express m-Cherry (red) and α-mCherry staining in AF647 (white). m-Cherry staining reveals >92% specificity for astrocytes (100 out of 105 neurons did not express m-Cherry) and penetrance of >99% (262 out of 264 astrocytes expressed m-cherry). Scale bars = 50μm. **(B)** For the CLARITY experiments, mice performance was enhanced during recent recall when CNO was administered at the moment of acquisition (CNO n=4 Saline n=2, p=0.003). **(C-D)** Number of tdTomato positive axons heading toward the NAc (p=0.95) **(C)** and the SuMB (p=0.23) **(D)** did not change when CNO was introduced during memory acquisition to mice expressing hM3Dq in CA1 astrocytes, compared to the control group. Data presented as mean ± SEM.

## Methods

### Mice

#### Ai14 mice

B6;129S6-Gt(ROSA)26Sor^tm14(CAG-tdTomato)Hze^, J-Stock No: 007908. These mice contain a loxP cassette with a stop codon, followed by CAG promoter-driven red fluorescent protein (tdTomato)(*68*).

#### TRAP mice

Fos^tm2·1(icre/ERT2)Luo^, J-Stock No: 030323. Knock-in mice with the CreER protein under the cFos promoter (*69*).

#### Wild type mice

C57BL/6J.

### Stereotactic Injections

Mice were anesthetized with isoflurane, and their heads were placed in a stereotactic apparatus (Kopf Instruments, USA). The skull was exposed and a small craniotomy was performed. Mice were bilaterally microinjected using the following coordinates – for the CA1: Anteroposterior (AP), −1.85mm from Bregma, Mediolateral (ML), ±1.4mm, Dorsoventral (DV), −1.5mm. For the ACC: AP +0.35mm, ML ±0.35mm, DV −1.8mm. For the NAc: AP −1.2mm, ML ±1.1mm, DV −4.5mm. All microinjections were performed using a 10μL syringe and a 34-gauge metal needle (WPI, Sarasota, USA). The injection volume and flow rate (0.1 ml/min) were controlled by an injection pump (WPI). Following each injection, the needle was left in place for 10 additional minutes to allow for diffusion of the viral vector away from the needle track and was then slowly withdrawn. The incision was closed using sewing and Vetbond tissue adhesive. For postoperative care, mice were subcutaneously injected with Tramadex (5mg/kg).

### Viral Vectors

AAV5-cFos::cre^ER^, AAVretro-CaMKII::iCre, AAVretro-CaMKII::GFP, AAV8-GFAP:: hM3D(Gq)-mCherry, AAV2-CAG::flex-TC66T-mCherry, AAV2-CAG::flex-oPBG were all from the ELSC Vector Core Facility. AAV5-hSyn::DIO-hM4Di-mCherry was purchased from Addgene (# 44362).

ENV-Rb-ΔG-eYFP was purchased from the viral vector core at the Kavli institute in Norway.

### Fear Conditioning

The fear conditioning apparatus consisted of a conditioning cage with a grid floor wired to a shock generator and surrounded by an acoustic chamber. To induce fear conditioning, mice were placed in the cage for two minutes, and a pure tone (2.9 kHz) was then sounded for 20 seconds followed by a two second foot shock (0.6mA). This procedure was then repeated, and 30 seconds after the delivery of the second shock, mice were extracted from the conditioning cage. Fear was assessed by a continuous measurement of freezing (complete immobility), the dominant behavioral fear response. To test contextual fear conditioning, mice were placed in the original conditioning cage, and freezing was measured for five minutes. Contextual fear conditioning recall was measured at day 1, day 2, day 4, day 15, day 28, and day 30, depending on the group to which the mouse was assigned.

### 4-hydroxytamoxifen (4-OHT)

For Ai14 mice, 3 mg of 4-OHT (Sigma H7904) were dissolved in 120μl of ethanol, and 480μl of sunflower oil were then added to create total volume of 600μl. For TRAP mice the 4-OHT was three time more concentrated. Each solution was used on the day of preparation. One hour after the relevant memory related task, 4-OHT solution was given to all mice (25mg/kg; i.p.) to enable CreER-mediated recombination.

### CNO

#### IP injections

CNO (Tocris #4936) was dissolved in DMSO and then diluted in 0.9% saline to yield a final DMSO concentration of 0.5%. Saline solution for control injections also consisted of 0.5% DMSO. 10mg/kg CNO was intraperitoneally injected 30min before the behavioral assays for the Gi pathway activation of the recent engram neurons, and 3mg/kg for the Gq pathway activation in astrocytes. The chosen doses of CNO did not induce any behavioral signs of seizure activity.

#### CNO in drinking water

For chronic inhibition of the CA1→ ACC, 70mg CNO were dissolved in 1ml of DMSO and added together with 10ml sucrose to 1L of water. A single IP injection of 10mg/kg CNO was administered immediately after recent recall, followed by the same concentration in their drinking water. The control group received the same ingredients (DMSO, sucrose) without the CNO. Drinking bottle was replaced every 24 hours to ensure 10mg/kg/day.

### Immunohistochemistry

90 minutes after the last memory task, mice were transcardially perfused with cold PBS followed by immediate removal of the brain into 4% paraformaldehyde (PFA) in phosphate-buffered saline (PBS). Brains were postfixed overnight at 4°C and cryoprotected in 30% sucrose in PBS. Brains were sectioned to a thickness of 50 μm using a sliding freezing microtome (Leica SM 2010R) and preserved in a cryoprotectant solution (25% glycerol and 30% ethylene glycol in PBS). Free-floating sections were washed in PBS, incubated for 1 hr in blocking solution (1% of bovine serum albumin, BSA, and 0.3% Triton X-100 in PBS). For cFos staining, the relevant brain slices were incubated for 7 days at 4°C with rabbit anti-cFos primary antibody (Synaptic system, #226003), and for the mCherry staining, slices were incubated overnight at 4°C with rabbit anti-mCherry primary antibody (Invitrogen, #PA5-34974). Sections were then washed with PBS and incubated for 2 hr at room temperature with secondary antibody (1:500, AF 647, donkey anti rabbit, #711-605-152, Jackson laboratory) in 1% BSA in PBS. Finally, sections were washed in PBS, incubated with 4,6-diamidino-2-phenylindole (DAPI; Sigma 1μg ml^-1^), and mounted on slides with Mounting Medium (Dako, #S3025)

### CLARITY

Full hemispheres were cleared based on a modified protocol(*41*) derived from that described by Ye et al. (2016)(*28*). Briefly, >12 weeks old mice were transcardially perfused with ice cold PBS followed by 4% PFA, brains were removed and kept in 4% PFA overnight at 4°C. Brains were then transferred to a 2% hydrogel solution (PBS with: 2% acrylamide, bio-rad #161-0140; 0.1% Bisacrylamide, bio-rad #161-0142; 0.25% VA-044 initiator, Wako, 011-19365; 4% PFA) for 48 hr. The samples were then degassed with N_2_ for 45 min and polymerized in 37°C for 3.5 hr. After degassing, the samples were cut at the mid-sagittal plane. The samples were then washed overnight in 200 mM NaOH-Boric buffer (sigma, #B7901) containing 8% sodium dodecyl sulfate (SDS) (sigma, #L3771), to remove PFA residuals. Samples were then stirred in a clearing solution (100 mM Tris-Boric buffer, biolab, #002009239100 with 8% SDS) at 37°C for 3–4 weeks. After the samples became transparent, they were washed with PBST (PBS with 0.5% tritonX100; ChemCruz, #sc-29112A) for 48 hr at 37°C with mild shaking (replacing the PBST every 24 hr) and for another 24 hr with new PBST 0.5% at RT. Finally, the samples were incubated in the refractive index matched solution CLARITY Specific Rapiclear (RI = 1.45; SunJin lab, #RCCS002) O/N at 37°C and for two more days at room temperature before imaging.

Transparent samples were embedded onto a slide surrounded by hot-glue walls. Thin coverslip glass was the placed over the brain from above, closing all exits. CLARITY specific RapiClear solution was then inserted into the chamber, covering the entire brain. Expanded protocol can be found in JoVE (*70*).

### Confocal Imaging

Confocal fluorescence images were acquired using an Olympus scanning laser microscope Fluoview FV1000 using 4X and 10X air objectives, 10X, and 20X water immersion objectives or 20X oil immersion objectives. Images were created by imaging between 30μm-5000μm in depth and reconstructing them using IMARIS 9.1.2 software (Bitplane, UK).

### Imaris analysis

Using the ‘spot’ feature on the Imaris software, (x,y) coordination were manually counted and extracted for all marked cells, DAPI included. To calculate the percentage of expected overlap between two groups out of a specific group, for each slice, the number of cells in the relevant group was divided by the number of the total amount of cells (i.e. DAPI), and multiplied by the same division for the second group in order to estimate the *percentage* expected overlap. This number was then multiplied by DAPI in order to predict the *number* of estimated overlap cells, and later divided by the number of counted cells of the first group in order to estimate the percentage of overlap cells out of the relevant group. Finally, the percentage of actual overlap cells were calculated by dividing the empirical number of overlap cells by the number of counted cells of the first group. This parameter was later divided by the expected overlap, allowing us to calculate the fold from expected measurement:

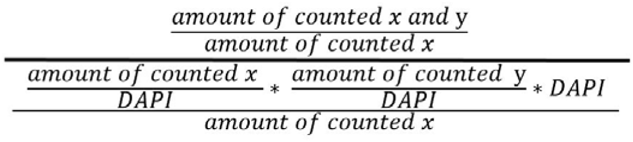

Where ‘*x*’ represents the sub group of cells from which we wanted to extract the percentage (i.e. ‘how many of x cells were also y cells, out of the total amount of x cells’).

### SyGlass analysis

Using the SyGlass virtual reality software, single axons were counted based on their destination inside visualized full hemispheres.

### Statistics

One-way Anova with Tukey post hoc and two tailed students t-test or paired t-test.

## Acknowledgements

We thank the entire Goshen lab for their support. This project has received funding from the European Research Council (ERC) under the European Union’s Horizon 2020 research and innovation program (grant agreement No 803589), the Israel Science Foundation (ISF grant No. 1815/18), and the Canada-Israel grants (CIHR-ISF, grant No. 2591/18). We thank Ami Citri, Mickey London, Adi Doron, Lior Naggan, and Yael Morose for the critical reading of the manuscript.

